# Individual Differences in the Connectivity of Left and Right Anterior Temporal Lobes Relate to Modality and Category Effects in Semantic Categorisation

**DOI:** 10.1101/2020.06.22.140509

**Authors:** Tirso RJ Gonzalez Alam, Katya Krieger-Redwood, Megan Evans, Grace E Rice, Jonathan Smallwood, Elizabeth Jefferies

**Affiliations:** Department of Psychology, University of York, YO10 5DD, UK; York Neuroimaging Centre, Innovation Way, Heslington, York YO10 5NY, UK; MRC Cognition and Brain Sciences Unit, University of Cambridge, 15 Chaucer Road, Cambridge, CB2 7EF

**Author notes:** Corresponding authors: Elizabeth Jefferies & Tirso RJ Gonzalez Alam. Address for correspondence: Department of Psychology, University of York, YO10 5DD, UK. Declarations of interest: none.

**Keywords:** Hemispheric Differences, Semantic Representations, Modality, Anterior Temporal Lobes, fMRI, Intrinsic connectivity

## Abstract

Contemporary neuroscientific accounts suggest that ventral anterior temporal lobe (ATL) regions act as a bilateral heteromodal semantic hub. However, research also shows graded functional differences between the hemispheres relating to linguistic versus non-linguistic semantic tasks and to knowledge of objects versus people. Individual differences in connectivity from bilateral ATL and between left and right ATL might therefore give rise to differences in function within this system. We investigated whether the relative strength of intrinsic connectivity from left and right ATL would relate to differences in performance on semantic tasks. We examined resting-state fMRI in 74 individuals and, in a separate session, examined semantic categorisation, manipulating stimulus type (famous faces versus landmarks) and modality of presentation (visual versus verbal). We found that people with greater connectivity between left and right ATL were more efficient at categorising landmarks, especially when these were presented visually. In addition, participants who showed stronger connectivity from right than left ATL to medial occipital cortex showed more efficient semantic categorisation of landmarks regardless of modality of presentation. These findings show that individual differences in the intrinsic connectivity of left and right ATL are associated with effects of category and modality in semantic categorisation. The results can be interpreted in terms of graded differences in the strengths of inputs from ‘spoke’ regions, such as regions of visual cortex, to a bilateral yet partially segregated semantic ‘hub’, encompassing left and right ATL.

## 1. Introduction

Semantic cognition allows us to understand the world around us – including the meaning of words, objects, locations and people (Lambon Ralph et al., 2017; Patterson et al., 2007). Conceptual representations that underpin semantic performance across input modalities (e.g., words and pictures) and across different tasks are thought to be supported by the bilateral ventral anterior temporal lobes (ATL; Binney et al., 2010; Rice et al., 2015a, 2015b). An influential account of ATL function suggests that this region forms a semantic “hub” drawing together different features represented within ‘spokes’ (capturing visual, valence, language and auditory inputs) to form heteromodal concepts (Patterson et al., 2007). This integration of different aspects of knowledge is thought to occur in a graded fashion, with the most heteromodal semantic responses in ventrolateral ATL (Lambon Ralph et al., 2017; Murphy et al., 2017; Visser et al., 2012). As semantic representations are thought to reflect interactions between the hub and spokes, individual differences in the way people represent and retrieve different types of concepts or categories may emerge from distinct patterns of hub and spoke connectivity.

Neuropsychological and neuroimaging studies suggest that semantic representation draws on both left and right ATL. Severe degradation of conceptual knowledge is seen following bilateral ATL atrophy in semantic dementia, while other aspects of cognition remain largely intact (Lambon Ralph et al., 2017; Patterson et al., 2007). These deficits are most pronounced for semantic tasks that probe specific-level knowledge – including knowledge of unique entities such as people, and highly-specific concepts, such as types of car (Rogers et al., 2015). In contrast, patients with unilateral lesions following resection for temporal lobe epilepsy have measurable yet much milder semantic deficits (Rice et al., 2018a). This might reflect functional compensation by the intact ATL (Jung and Lambon Ralph, 2016). Neuroimaging studies with healthy participants have also found bilateral responses to semantic tasks in ATL (Visser et al., 2009), irrespective of whether words or pictures are presented (Bright et al., 2004; Tranel et al., 2005; Vandenberghe et al., 1996; Visser et al., 2012; Visser and Lambon Ralph, 2011). Moreover, inhibitory transcranial magnetic stimulation (TMS) delivered to either left or right ATL disrupts both picture and word-based semantic tasks, mimicking the pattern in semantic dementia (Pobric et al., 2010, 2007). In line with expectations for a single semantic hub distributed across two hemispheres (cf. Schapiro et al., 2013), inhibitory TMS to left ATL leads to an increase in the response within right ATL, suggesting the non-stimulated hemisphere may compensate for functional disruption within the stimulated hemisphere (Binney and Lambon Ralph, 2015; Jung and Lambon Ralph, 2016).

Nevertheless, there is also strong evidence that the left and right ATL are not functionally identical. There are differences in the extent to which the two ATLs show white matter connectivity to visual, auditory-motor, social and emotional networks (Papinutto et al., 2016), potentially resulting in a degree of functional specialisation. By this view, ventral ATL acts as a bilateral hub that integrates lower-level spokes in a graded fashion, with this multi-modal integration not occurring in an all-or-none fashion but instead reflecting gradients of connectivity within and between the ATLs to specific hubs (Bajada et al., 2019; Binney et al., 2012; Rice et al., 2015a). Some studies have suggested a modality difference – with left ATL showing stronger engagement for verbal tasks, and right ATL showing a preference for non-verbal tasks. This suggestion has some support from studies of patients with semantic dementia who have relatively more left-sided or right-sided atrophy (Gainotti, 2012). For example, Snowden et al. (2004) found that patients with more left-lateralised atrophy had greater impairment for people’s names, while patients with more right-sided atrophy had greater difficulty on semantic tasks employing faces. Similarly, several studies have shown that atrophy in right ATL correlates with difficulties on picture semantic tasks, while damage to left ATL is more strongly correlated with verbal semantic task performance (Butler et al., 2009; Mion et al., 2010). A variant of this modality view suggests that output modality is also important – damage to left ATL is associated with problems in naming concepts (Lambon Ralph et al., 2001), and therefore with deficient lexical access from semantic knowledge, while right ATL is linked to poor object recognition (Damasio et al., 2004).

An alternative account of functional specialisation across left and right ATL suggests it is not the input/output modality that is critical, but instead the nature of the conceptual information itself – by this view the right ATL has been argued to play a larger role than left ATL in understanding social concepts and retrieving conceptual information about specific people (Olson et al., 2013, 2007; Ross and Olson, 2010; Zahn et al., 2007). Patients with damage to right ATL often have difficulties recognising faces, but there is an ongoing debate about whether these difficulties reflect impairment for faces per se (i.e. difficulty when the task involves both social stimuli and picture inputs) or a wider problem with social concepts (Gainotti, 2013; Gorno-Tempini et al., 1998). A recent fMRI study (Rice et al., 2018b) directly compared the neural response in ATL during semantic decisions about specific entities that were social (people) and non-social (landmarks). The social and non-social stimuli were presented as both words (i.e., people’s names) and as pictures (i.e., of faces). This study found that both the left and right ventral ATL responded regardless of the modality or category of semantic information. However, an additional region in the ATL, extending towards the temporal pole also showed stronger activation to people vs. landmarks.

A recent neuroimaging meta-analysis of 97 functional neuroimaging studies (Rice et al., 2015b) confirmed the complex pattern of functional similarities and differences between left and right ATL. Both left and right ATL were activated across verbal and non-verbal stimuli, and social and non-social tasks. However, activation likelihood estimation revealed that studies involving word retrieval were more likely to report unilateral left ATL activation, while social semantic studies were more likely to observe bilateral ATL activation (with non-social tasks equally likely to be either bilateral or left lateralised, since many of these tasks involved naming). This functional complexity is also reflected in the dorsoventral axis of ATL: dorsal aspects show greater intrinsic connectivity with auditory/language-related areas, while posterior ventral aspects show greater coupling with visual networks (Jackson et al., 2017; Murphy et al., 2017). Converging fMRI, TMS and neuropsychological evidence shows an important role of dorsal ATL in social processing, with more right than left superior ATL involvement for social compared to non-social matched stimuli (Binney et al., 2016; Pobric et al., 2016), and with left superior ATL playing a necessary role in the retrieval of both social and non-social abstract concepts. A similar distinction has been reported for ventral ATL, with words eliciting left ventral ATL activation, while objects engage bilateral ventral ATL, although a right bias for objects does appear in more posterior temporal cortex (Hoffman and Lambon Ralph, 2018). Given that Rice and colleagues suggested these functional differences reflect differential connectivity between left and right ATL and other brain networks – for example, stronger connectivity between right ATL and regions associated with social cognition, or between left ATL and language regions – we might anticipate that individual differences in intrinsic connectivity at rest would relate to performance on social vs. non-social, or verbal vs. non-verbal semantic tasks.

In a recent study (Gonzalez Alam et al., 2019), we compared the intrinsic connectivity of four heteromodal semantic sites across hemispheres (ATL, angular gyrus, posterior middle temporal gyrus and left inferior frontal gyrus), and found that ATL had the most symmetrical connections (i.e. the highest correlations between connectivity patterns generated from left and right-hemisphere seeds). However, some subtle differences in connectivity were still observed. Left ATL was more connected with other sites implicated in semantic cognition, including left inferior frontal gyrus, posterior middle and inferior temporal cortex, posterior dorsal angular gyrus/intraparietal sulcus and medial temporal lobe. Right ATL was more connected to visual cortex, as well as to extensive regions of default mode network, including angular gyrus and dorsomedial prefrontal cortex. Individual differences in these patterns of connectivity from left and right ATL might be expected to differentially affect the efficiency of semantic decisions about specific social and non-social concepts, presented as words and pictures. Although Gonzalez Alam et al. (2019) failed to observe any behavioural correlates of hemispheric differences in ATL connectivity, the tasks used in this previous study were not designed to maximise the involvement of this region, as they did not use specific-level concepts, or compare social and non-social concepts. To address this issue, the current study acquired resting-state fMRI from 74 participants, who completed semantic decisions about specific social and non-social concepts, presented as written words and pictures (using the stimuli from Rice et al., 2018b) in a separate testing session following the scan. We then assessed relationships between connectivity and behavioural performance – with particular focus on whether right vs. left ATL connectivity would predict performance on social vs. non-social tasks and trials presented as pictures vs. words.

## 2. Methods

### 2.1. Participants

This study was approved by the local research ethics committees. Participants who had previously received a resting-state scan were invited to return to the lab for behavioural testing. The recruited sample consisted of 83 participants (19 male, 64 female, mean age=19.69, range=18-26). Two participants were removed before pre-processing due to missing resting-state scans, and a further three due to not having full brain coverage. Another two were excluded during pre-processing because they exceeded our quality assessment measures of (i) motion greater than 0.3mm, (ii) invalid scans greater than 20%; and/or (iii) global mean signal change greater than z=2. Finally, two more participants were excluded because they performed at least one behavioural task at chance level, leaving us with a final sample size of 74 participants recruited from undergraduate and postgraduate students at the University of York. All participants were right handed, native English speakers with normal/corrected vision. During scanning, we also excluded participants with a history of psychiatric or neurological illness, severe claustrophobia, drug use that could alter cognitive functioning, and pregnancy. All volunteers provided written informed consent and were either paid or given course credit for their participation.

### 2.2. Procedure

In the initial neuroimaging session, we acquired structural MRI images and a resting-state scan during which participants were instructed to keep their eyes open and focus on a fixation cross. We invited participants back for a behavioural session involving five tasks. The duration of the full testing session was approximately 1.5 hours. The task used in the current study was towards the end of this session. To control for order effects, each participant completed the tasks in the same order.

### 2.3. Task

We adapted a task from Rice et al. (2018b). Participants were presented with different categories of stimuli (animals, landmarks and people) as either images or written words (in the original task the latter were presented as spoken words). They had to judge whether the stimuli were European or non-European. They were also presented with a non-semantic perceptual control condition, in which participants were shown a scrambled image (generated by taking the pixels from the images in the other conditions and randomising their location so they were devoid of meaning) and asked to judge whether it was presented higher or lower on the screen. Examples of the stimuli in each condition are shown in Figure 1.

**Figure 1.**
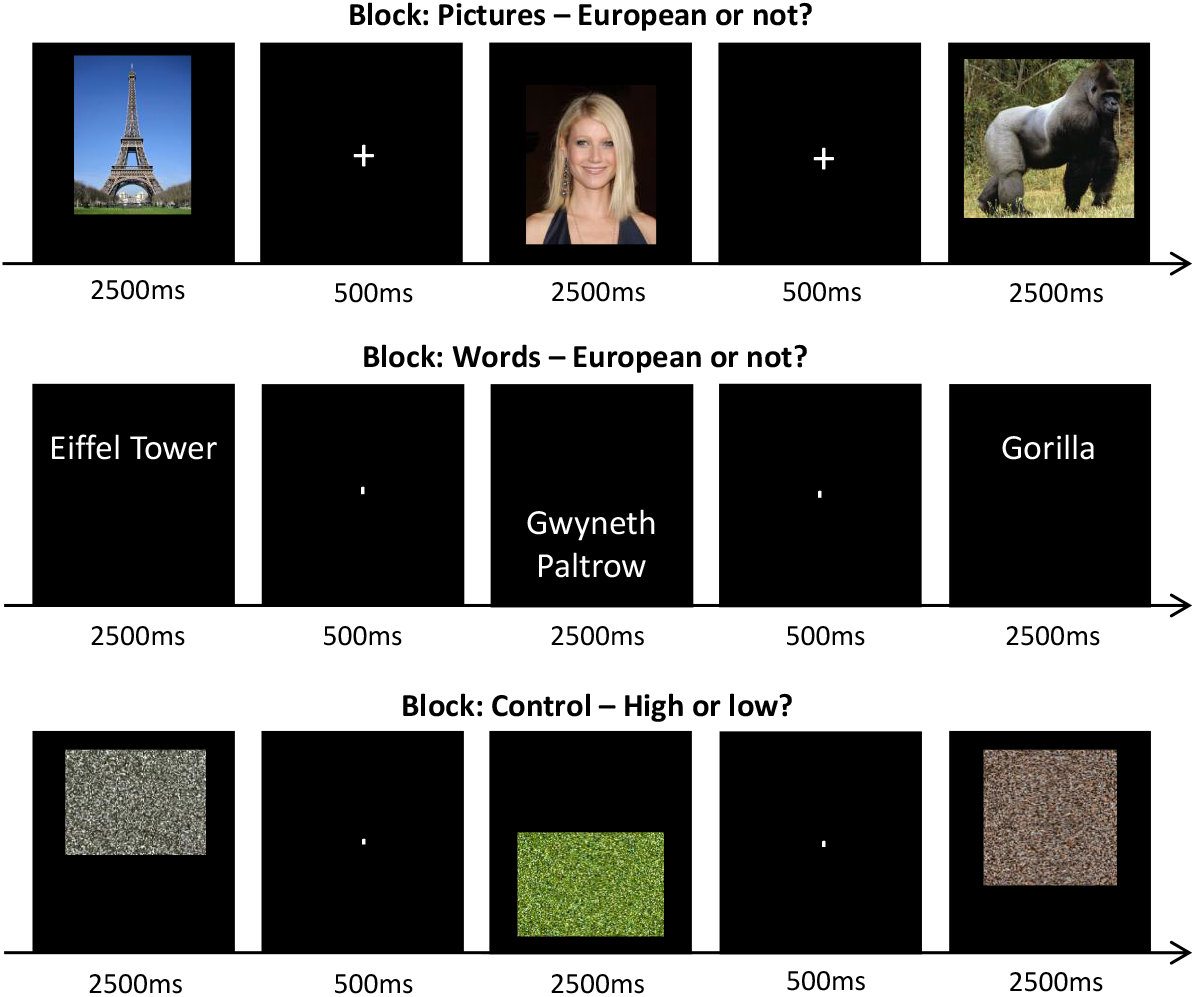
Example stimuli and trial structure for each condition in the semantic representation task and non-semantic control task. This is a simplification of the actual structure of the task, where the stimuli were not only blocked by modality of presentation, but also by category of stimuli (i.e. “pictures of landmarks” would be a block).

Stimuli were taken from Rice et al. (2018b) and reduced to only include trials with 85% accuracy or above in Rice’s data, providing 60 stimuli per category. Blocks consisted of 30 trials each (480 trials in total), all drawn from the same category and modality, with random assignment of stimuli to blocks. There were sixteen blocks (four conditions by two modalities; with each combination presented twice). The blocks were presented in four possible sequences, to which participants were randomly assigned.

Each trial lasted 3s, consisting of a 500ms fixation cross followed by 2500ms stimulus presentation (see Figure 1). Participants indicated their response using the ‘1’ key on a computer keyboard for European/higher location and ‘2’ for non-European/lower location. Before the task commenced, an instruction slide was presented which remained on screen until the participant indicated they were ready to begin via key press. At the beginning of each block, a cue screen indicating the condition was presented for 500ms. Every four blocks participants were presented with a rest screen until they indicated they were ready to continue via key press. Both response time (RT) and accuracy were recorded, and an efficiency score was calculated for each participant in each condition by dividing median response times by accuracy (note: in brain analyses, this efficiency score was inverted to aid the interpretation of the results, such that a higher score corresponded to better performance). The duration of the whole task was 15-20 minutes. The task was implemented in E-prime 2.0.

### 2.4. Neuroimaging

#### 2.4.1. MRI data acquisition

MRI data was acquired using a 3T GE HDx Excite Magnetic Resonance Imaging (MRI) system utilising an eight-channel phased array head coil tuned to 127.4 MHz, at the York Neuroimaging Centre, University of York. Structural MRI acquisition in all participants was based on a T1-weighted 3D fast spoiled gradient echo sequence (TR = 7.8s, TE = minimum full, flip angle = 20°, matrix size = 256 × 256, 176 slices, voxel size = 1.13 × 1.13 × 1 mm). A nine-minute resting state fMRI scan was carried out using single-shot 2D gradient-echo-planar imaging (TR = 3s, TE = minimum full, flip angle = 90°, matrix size = 64 × 64, 60 slices, voxel size = 3 × 3 × 3 mm^3^, 180 volumes). Participants were asked to passively view a fixation cross and not to think of anything in particular during the resting-state scan. A FLAIR scan with the same orientation as the functional scans was collected to improve co-registration between subject-specific structural and functional scans.

#### 2.4.2. Pre-Processing

All pre-processing of resting-state data was performed using the CONN functional connectivity toolbox V.18a (http://www.nitrc.org/projects/conn; Whitfield-Gabrieli & Nieto-Castanon, 2012). MRI data pre-processing and statistical analyses were carried out using the SPM software package (Version 12.0), based on the MATLAB platform (Version 17a) implemented in CONN. For pre-processing, functional volumes were slice-time (bottom-up, interleaved) and motion-corrected, skull-stripped and co-registered to the high-resolution structural image, spatially normalized to the Montreal Neurological Institute (MNI) space using the unified-segmentation algorithm, smoothed with a 6 mm FWHM Gaussian kernel, and band-passed filtered (0.008 – 0.09 Hz) to reduce low-frequency drift and noise effects. A pre-processing pipeline of nuisance regression included motion (twelve parameters: the six translation and rotation parameters and their temporal derivatives), scrubbing (all outlier volumes were identified through the artifact detection algorithm included in CONN, with conservative settings: scans for each participant were flagged as outliers based on a composite metric with parameters set to scan-by-scan change in global signal z-value threshold = 3, subject motion threshold = 5mm, differential motion and composite motion exceeding 95% percentile in the normative sample) and CompCor components (the first five) attributable to the signal from white matter and CSF (Behzadi et al., 2007), as well as a linear detrending term, eliminating the need for global signal normalization (Chai et al., 2012; Murphy et al., 2009).

#### 2.4.3. ROI Selection

We used a left ventral ATL site identified by Rice et al. (2018b) as showing peak activation for semantic > non-semantic contrasts across eight studies using distortion-corrected fMRI (MNI coordinates −41, −15, −31). Since there is good evidence of bilateral engagement of ATL in semantic cognition, and Rice et al. (2018b) identified a right ATL functional peak (MNI 44, −11, −36), we also included this seed in our investigation. We created ROIs for both sides by placing a binarised spherical mask with a radius of 3mm at the MNI coordinates identified above. The BOLD time series extracted for each seed region was an average for all voxels making up the sphere.

#### 2.4.4. Resting-State fMRI Analysis

This analysis examined individual differences in the connectivity of left and right ATL to the rest of the brain, measured through resting-state fMRI, and related these differences to behavioural efficiency scores on semantic tasks (measured outside the scanner in a separate session). In a first-level analysis, we extracted the time series from each of the two ROIs for each participant. We computed the seed to voxel correlations for each of our seeds, removing the nuisance regressors detailed in section 2.4.2. At the group level, our analysis focused on associations between hemispheric similarities and/or differences in the connectivity of ATL and behavioural effects of the category of the stimuli and the modality of presentation. We entered into a GLM the mean-centred efficiency scores (with outliers +/-2.5 SD imputed to +/-2.5) of five task conditions (excluding the Animal Verbal condition, and the non-semantic Control condition, due to performance levels at chance and at ceiling, respectively), together with a nuisance regressor containing mean motion (measured in framewise displacement) for each participant as EVs.

We performed functional connectivity weighted GLM seed-to-voxel analyses convolved with a canonical haemodynamic response function (HRF). There were four analyses: two models for the single seeds (left ATL, right ATL); one model examining the common connectivity of left and right ATL (examining seed regions that encompassed both hemispheres); and finally one model that examined the difference between left and right ATL (examining left versus right functional peaks). We applied Bonferroni correction to the FWE values resulting from two-sided t-tests in the models to determine significant clusters (correcting for these four models). Besides the mean group connectivity for each seed, we defined the following contrasts of interest: category (People > Landmarks and vice-versa), modality (Verbal > Visual and vice-versa), modality by category interaction (the effect of modality for people versus landmarks) and the main effects for the five tasks conditions (i.e. Verbal Landmark, Verbal People, Visual Landmark, Visual People and Visual Animals). The single seed models did not yield significant results and are therefore not discussed further.

At the group-level, analyses were carried out using CONN with cluster correction (with results reaching p < .013 considered to be significant, corresponding to p < .05 following Bonferroni correction to control for the four models we ran), and a threshold of z=3.1 (p-FWE=0.001) to define contiguous clusters (Eklund et al., 2016). This analysis included the behavioural regressors described above (as mean-centred inverse efficiency scores for each task condition) to evaluate whether performance correlated with individual differences in intrinsic connectivity. The connectivity maps resulting from these analyses were uploaded to Neurovault (Gorgolewski et al., 2015, URL: https://neurovault.org/collections/5687/). As a confirmatory analysis, to verify that the results were not dependent on an arbitrary significance threshold, we carried out non-parametric permutation testing as implemented in CONN for each significant result that survived Bonferroni correction. All clusters were replicated by permutation testing.

In order to interpret the results that survived Bonferroni correction, we used the significant clusters as binarised masks and extracted the global-scaled mean connectivity for each seed per participant to each significant cluster using REX implemented in CONN (Whitfield-Gabrieli & Nieto-Castanon, 2012). These values were then related to each participant’s mean-centred inverse efficiency score for the relevant EV, and plotted as scatterplots using Seaborn in Python 2.7, colour coded so that red scatterplots show connectivity from the left ATL seed and blue scatterplots show connectivity from the right ATL seeds.

#### 2.4.5. Control Analysis

Due to the functional heterogeneity of ATL (Bajada et al., 2019; Binney et al., 2012; Rice et al., 2015a), the exact seed location can have an impact on the results. Since in a previous study we found significantly different patterns of cross-hemisphere connectivity in ATL depending on whether sign-flipping or functional peaks were used to define the right-hemisphere seed (Gonzalez Alam et al., 2019), we carried out a control analysis using sign-flipping to determine homotopic left and right ATL. This analysis assessed the robustness of the findings obtained from the functionally-defined right ATL seed.

For this supplementary control analysis, we generated a right-hemisphere sphere using coordinates for the left ventral ATL peak identified by Rice et al. (2018b), and flipping the sign of the x-coordinate in MNI space from negative to positive (MNI coordinates 41, −15, −31). In this control analysis, left and right ATL sites were anatomically equivalent, while in our main analysis, we used functional peaks in left and right ATL which were in similar but non-identical locations (see supplementary figure S3 for the relative position of the right ATL seeds). We created a ROI by placing a binarised spherical mask with a radius of 3mm at the MNI coordinates identified above and replicated the main analysis pipeline, with the exception of the Bonferroni correction, which was set to correct for 7 multiple comparisons to account for the three extra supplementary analyses (one single RH seed and both its conjunction and difference with LH). We performed the same GLM with this homotopic sign-flipped right-hemisphere seed as with the functionally-defined seed to guard against Type 1 errors. We also binarised the significant results obtained from the functionally-defined seed in our main analysis below and examined whether the same patterns could be observed using the homotopic sign-flipped seeds, to assess the robustness of our main findings. For these supplementary analyses, we extracted the connectivity of the left functional and right sign-flipped seeds, and regressed these patterns of connectivity against the relevant behavioural contrast that gave rise to each neuroimaging effect. This allowed us to verify whether the effects we report below were dependent on seed selection or could be recovered regardless of the method used to define homotopy. All of the results reported below were significant regardless of the method used to define homotopy. The detailed methods and results of this control analysis are provided in the supplementary materials.

## 3. Results

### 3.1. Behavioural Results

Figure 2 shows the average median reaction time, accuracy and efficiency scores in each condition of the semantic task adapted from Rice et al. (2018b). A two-way repeated measures ANOVA on reaction time data with category and modality as factors revealed no main effect of modality, a significant main effect of category and a category by modality interaction [Category: F(2,144) = 56.07, p < .001; Interaction: F(2,144)=69.1, p < .001]. Post-hoc tests with Bonferroni correction showed no difference between people and landmarks, but significant differences between animals and all other conditions, with participants responding more slowly on animal trials (p < .001). Post-hoc testing with Bonferroni correction also confirmed the interaction was driven by slower performance for verbal than visual animal judgements (p < .001). The same analysis for accuracy found significant effects of category, modality and a category by modality interaction [Category: F(2,144) = 174.39, p < .001; Modality: F(2,144) = 487.39, p < .001; Interaction: F(2,144) = 41116.11, p < .001]. Post-hoc tests with Bonferroni correction revealed that participants showed equivalent accuracy for landmark and people judgements, but both of these significantly differed from animal judgements, where participants made more errors (p < .001). Likewise, participants were significantly less accurate in verbal than picture judgements (p < .001). The interaction was driven by participants being significantly less accurate for verbal than visual judgements about animals (p < .001).

**Figure 2.**
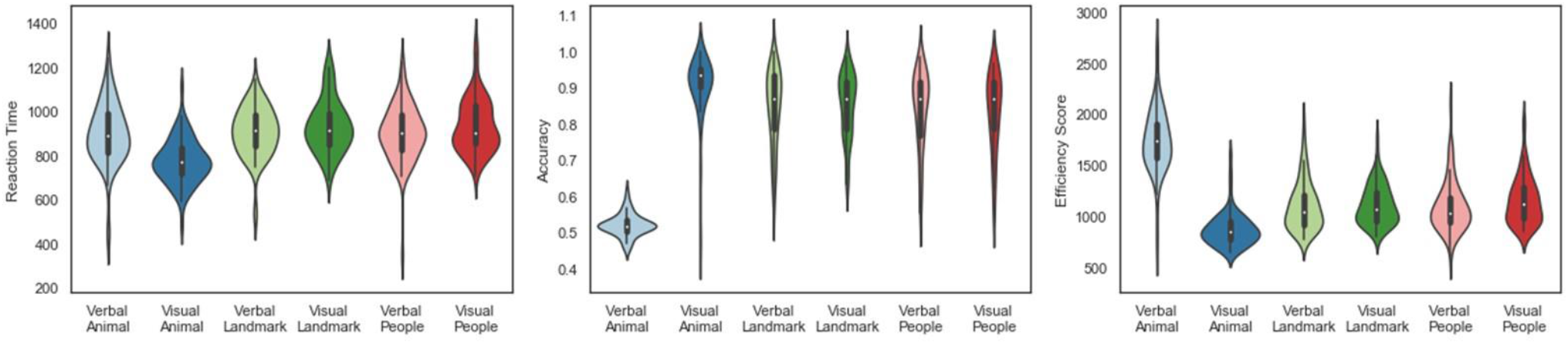
Violin plots depicting the average median reaction time (milliseconds), accuracy (proportion correct) and efficiency scores (reaction time divided by accuracy) for the semantic categorisation task. The width of each bar shows the frequency of scores for the values plotted.

Speed and accuracy may be traded off in different ways across tasks and individuals. To address this issue we calculated response efficiency scores, computing reaction time divided by accuracy to characterise global performance (inverted for neuroimaging analyses, so that high scores reflect good performance in the scatterplots depicted in Figures 4 – 6 and supplementary figures: these figures depict each participant’s efficiency z-scored relative to the group performance in each condition). A two-way repeated measures ANOVA of participants’ efficiency scores with Greenhouse-Geisser correction using category and modality as factors showed significant effects of category, modality and an interaction using this response efficiency measure [Category: F(1.7,122.6) = 63.22, p < .001; Modality: F(1,72) = 136.08, p < .001; Interaction: F(1.8,130)=478.47, p < .001]. Post-hoc tests once again found no difference between judgements about landmarks and people, plus significantly better performance in both of these conditions compared with animal trials (p < .001). Participants performed less efficiently in verbal than visual judgements (p < .001), while the interaction was driven by participants being less efficient in verbal than visual judgements about animals (p < .001).

### 3.2. Mean connectivity of left and right ATL (without behavioural regressors)

We examined the single seed mean connectivity for each ATL, as well as mean connectivity from both seeds combined, and differential left vs. right connectivity. The results are shown in Figure 3. The left ATL seed showed a pattern of bilateral functional connectivity overlapping with the semantic cognition network (shown for reference in the top row) in the LH, including medial and lateral aspects of the temporal lobe extending posteriorly to the angular gyrus bilaterally, as well as parts of the inferior frontal gyrus (extending more anteriorly in the LH) and central sulcus, posterior cingulate cortex and frontal poles (Figure 3, second row). The right ATL seed showed positive connectivity to ventral aspects of the left temporal lobe and right temporal pole. It also showed positive connectivity to left angular gyrus, inferior frontal and posterior middle temporal gyri, frontal pole and medial aspects of the temporal lobe, plus weak connectivity to bilateral occipital lobes. Unlike left ATL, the RH seed did not show connectivity to the central sulcus, but it did to bilateral superior frontal gyrus and medial orbitofrontal cortex extending posteriorly to the posterior cingulate cortex (Figure 3, third row). This pattern is similar to that described by Gonzalez Alam et al. (2019). An analysis giving equal weight to both left and right ATL seeds captured this similarity between the maps, including strong connectivity in bilateral temporal regions extending into angular gyrus, as well as inferior and superior frontal gyri, posterior cingulate and orbitofrontal cortex, with common weak connectivity in medial occipital cortex (Figure 3, fourth row). Significant differences between the left and right ATL included the left ATL seed showing stronger connectivity with left dorsal ATL and bilateral dorsal central sulcus, while the right ATL seed showed stronger connectivity with left middle frontal gyrus, orbitofrontal and ATL (Figure 3, bottom row).

**Figure 3.**
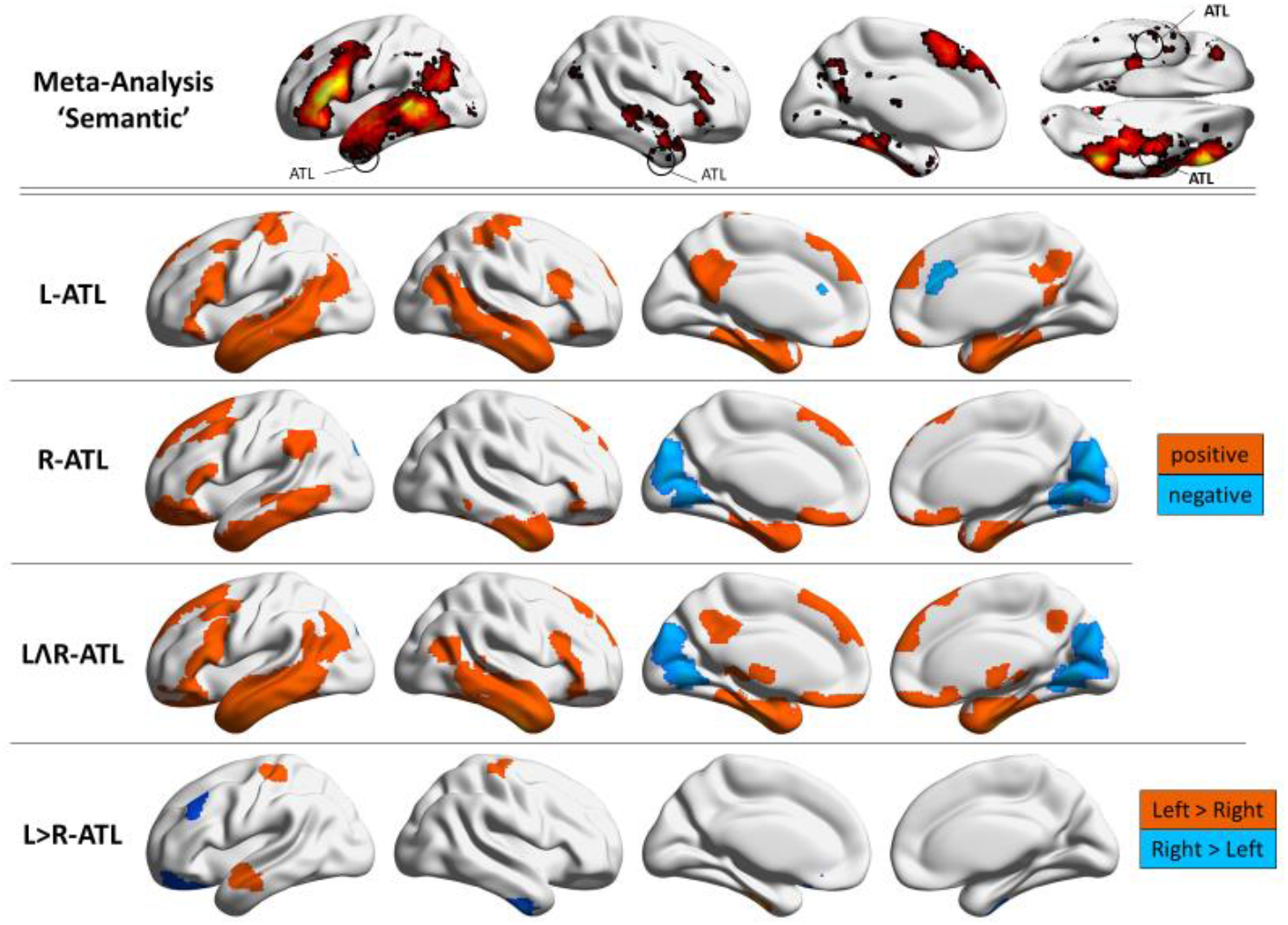
The top row depicts a term-based meta-analysis of the term ‘Semantic’ in Neurosynth. The four following rows show resting-state connectivity for left and right anterior temporal lobe, plus their common and differential connectivity. For the single seeds and mean left plus right ATL maps, the warm and cool colours represent positive and negative patterns of connectivity respectively, while for the difference analysis the warm and cool colours represent left and right connectivity respectively. The group maps are thresholded at z = 3.1, p = 0.05.

### 3.3. Behavioural consequences of ATL connectivity – overview of analysis aims

We performed whole brain resting-state functional connectivity analysis using behavioural performance in five conditions of our semantic representation task as covariates, to probe for possible associations between seed connectivity and categorisation efficiency. We did not include the verbal animal condition due to chance-level performance. For each significant result found using a cluster-forming threshold of z = 3.1, we ran non-parametric analysis using CONN to confirm whether the result was robust irrespective of this particular cluster-forming threshold. All results presented in this section, except for the occipital pole effect, were replicated across these analyses. Table 1 presents a full list of significant results, after Bonferroni correction for 4 comparisons, as well as which results are significant using non-parametric statistics. In the following sections, we first present the results of analyses for combined left and right ATL seeds (i.e. mean connectivity across both hemispheres), followed by analyses of differences between left and right ATL connectivity. Behavioural associations for the single seeds (i.e. for left and right ATL seeded separately) were captured by the results of the combined analyses reported below and consequently these findings are not discussed below although they are reported full in the supplementary materials.

**Table 1.** Summary of significant results. Note. The p-FWE values are Bonferroni-corrected for four multiple comparisons. The “Non-param. p” column reports the significance value (p-FWE) obtained using non-parametric statistics (5,000 simulations) as implemented in CONN. L = Left; R = Right; ATL = Anterior Temporal Lobe; TP = Temporal Pole; OP = Occipital Pole

| Seed | Contrast | Cluster | Vox. | p-FWE | Non-param. p | x | y | z |
| --- | --- | --- | --- | --- | --- | --- | --- | --- |
| Single Seeds |  |  |  |  |  |  |  |  |
| L-ATL | Visual > Verbal<br>Landmark | R-ATL | 242 | <.001 | .035 | 48 | 10 | -44 |
| L-ATL | Interaction | R-ATL | 169 | .032 | >.05 | 40 | 14 | -46 |
| R-ATL | Visual > Verbal<br>Landmark | L-TP | 575 | <.001 | .003 | -42 | -4 | -34 |
|  |  | R-TP | 175 | .012 | >.05 | 38 | 12 | -42 |
| R-ATL | Visual Landmark | R-TP | 158 | .042 | >.05 | 38 | 12 | -42 |
| Combined Results |  |  |  |  |  |  |  |  |
| L and R ATL | Interaction | R-TP | 186 | .02 | .047 | 48 | 6 | -40 |
| L and R ATL | Visual Landmark | R-TP | 292 | <.001 | .02 | 48 | 10 | -36 |
| L and R ATL | Visual > Verbal<br>Landmark | L-ATL | 649 | <.001 | .002 | -42 | -4 | -48 |
| L and R ATL |  | R-ATL | 539 | <.001 | .004 | 38 | 12 | -44 |
| R > L ATL | Main Landmark | L-OP | 187 | .016 | > .05 | -12 | -100 | 8 |

The results converged on two main findings: (i) the strength of connectivity between right ATL and medial occipital cortex is associated with efficient performance on landmarks trials, regardless of modality; and (ii), the strength of connectivity between left and right ATLs (especially stronger bilateral ATL connectivity) was linked to better performance on landmark trials in the picture modality. We show the overlap of different results relating to this latter effect in Figure 7.

### 3.4 Mean connectivity across left and right ATL and associations with behaviour

The mean connectivity of left and right ATL was significantly associated with three behavioural results. First, we found a significant interaction between modality and category (see Figure 4). Stronger connectivity from the combined left and right ATL seed to a more anterior right-lateralised cluster in the temporal pole was associated with better performance in the visual than verbal modality for landmarks, whilst the reverse pattern was observed for trials involving people knowledge (relatively better performance in the verbal than visual modality). We also found two significant results for the landmarks task, consistent with this interaction. Connectivity of the bilateral seed to a bilateral cluster encompassing left and right ATL, extending to the temporal pole, was associated with better visual than verbal categorisation for landmarks (see Figure 5, top panel). People who were better at categorising visual landmarks overall also had stronger connectivity between the bilateral ATL seed and right temporal pole (Figure 5, bottom panel). In order to confirm that these findings reflected cross-hemispheric connectivity, we plotted the results for ATL seed regions within left and right hemisphere separately. These plots show that left and right-lateralised parts of the seed were both associated with better categorisation for the picture landmark condition. All of these results taken together suggest that stronger connectivity to bilateral temporal pole is seen in participants who show better categorisation of landmarks when these are presented as pictures as opposed to words.

**Figure 4.**
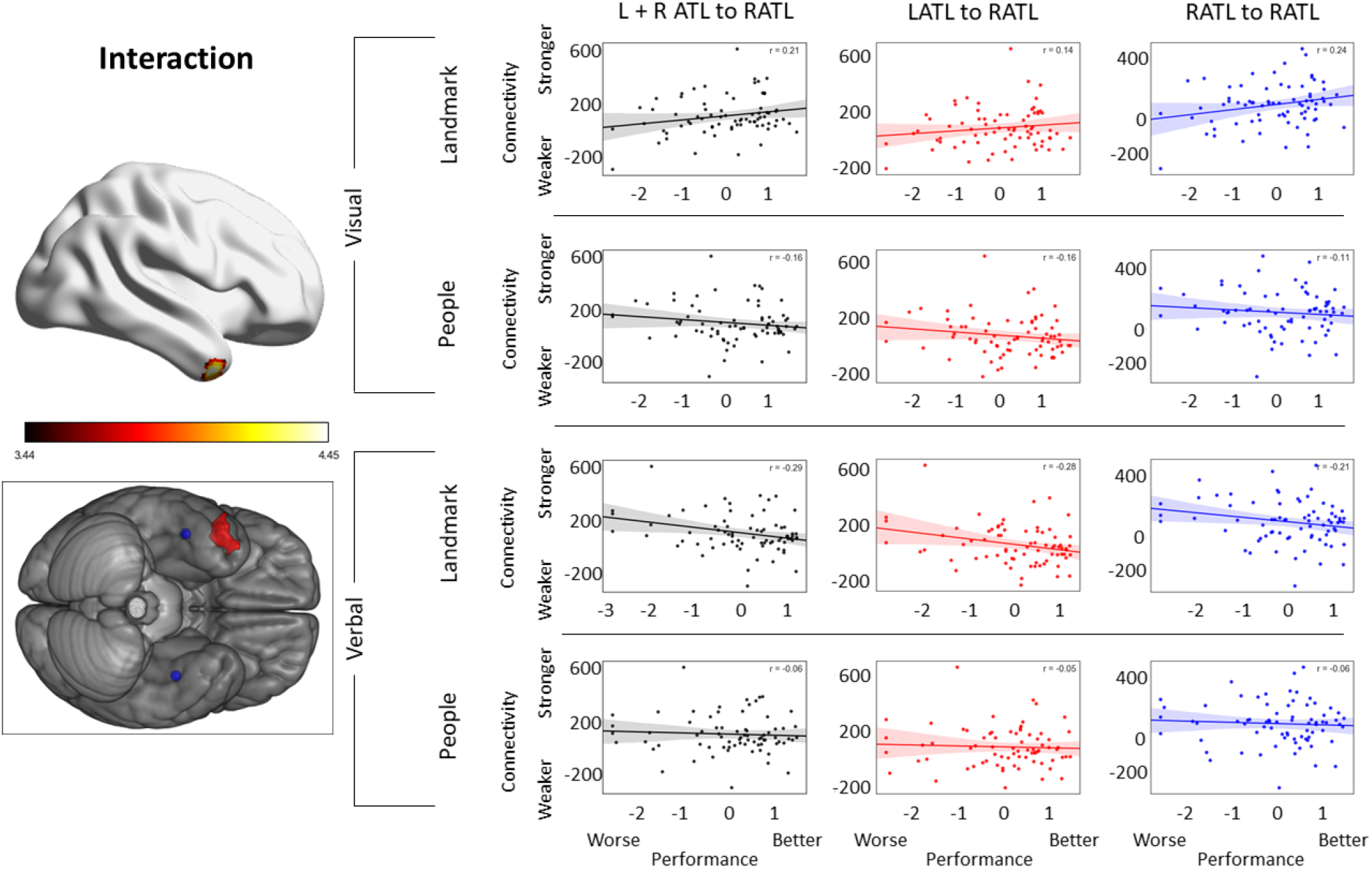
The connectivity between a right temporal pole cluster and the bilateral ventral ATL seed (defined using distinct functional peaks in both hemispheres) was associated with a modality by category interaction (Z=3.1, p=.05, Bonferroni-corrected for 4 models). The scatterplots depict the mean-centred efficiency scores (given in z-scores), plotted as a function of the normalised functional connectivity (i.e. scaled to the global mean) between the right temporal pole cluster depicted in the figure and seed regions comprising both left and right ATL functionally-defined seeds (black), left functionally-defined ATL (red), and right functionally-defined ATL (blue). The seeds have been plotted alongside the cluster on a separate brain image to depict their relative position. All units are given in z-scores.

**Figure 5.**
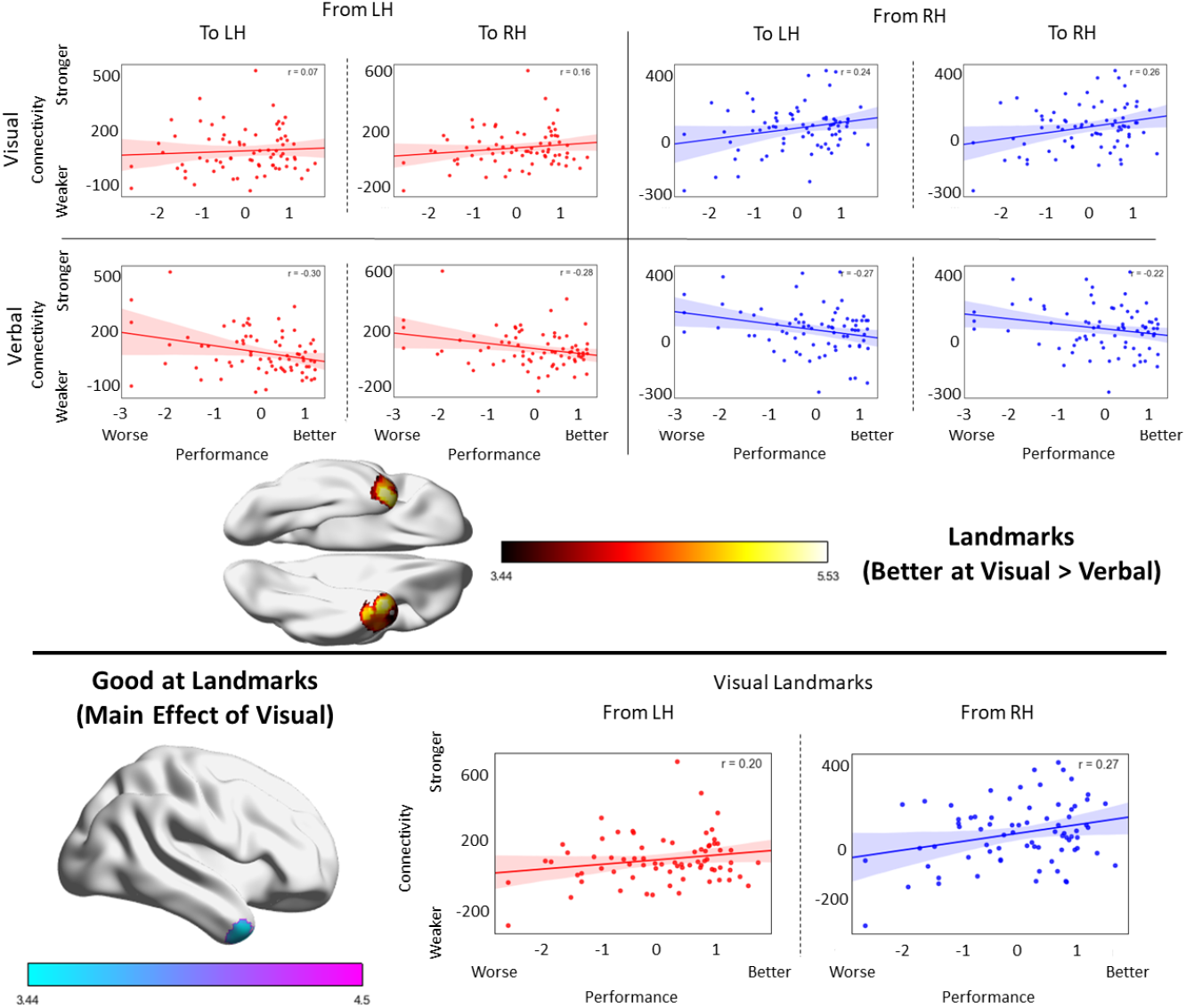
Top panel: Bilateral temporal pole clusters whose connectivity to bilateral left and right ventral ATL was significantly associated with being better at visual relative to verbal landmark judgements. Bottom panel: a right temporal pole cluster whose connectivity to bilateral left and right ventral ATL was significantly associated with the main effect of being good at visual landmarks (both results are thresholded at Z=3.1, p=.05, Bonferroni-corrected for 4 models). The scatterplots depict the mean-centred efficiency scores in the relevant condition (given in z-scores) plotted as a function of the normalised functional connectivity (i.e. scaled to the global mean) from both left ATL (red) and right ATL (blue) functional peaks to the clusters depicted in the figure. All units are given in z-scores.

### 3.5. Differential ATL connectivity between hemispheres and associations with behaviour

In a second set of analyses, we examined whether differences in connectivity between left and right ATL related to semantic performance. A whole-brain difference analysis contrasting left ATL over right ATL revealed a significant result located in medial occipital lobe / occipital pole, relating to performance for landmarks, regardless of modality (Figure 6). Participants with more right than left ATL connectivity to this occipital pole cluster showed better performance for landmarks.

**Figure 6.**
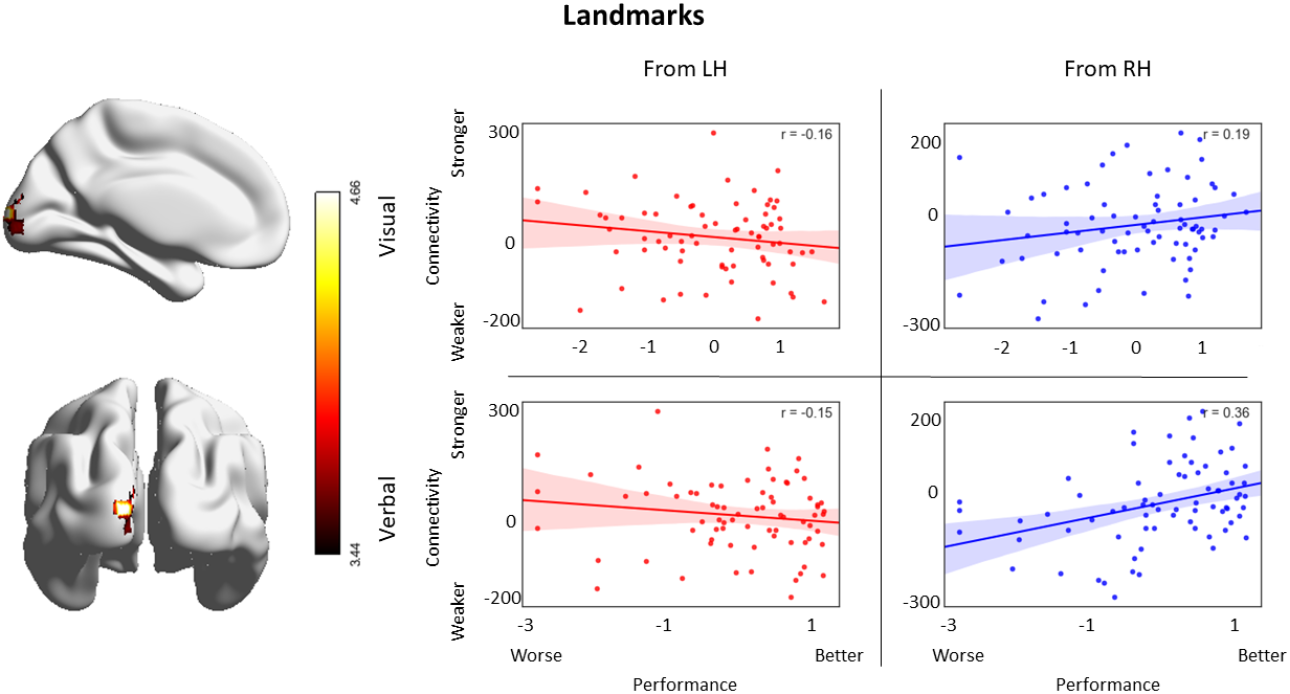
A polar occipital cluster whose differential connectivity to left ATL versus functionally-defined right ATL was significantly associated with landmarks performance (Z=3.1, p=.05, Bonferroni-corrected for 7 models). The scatterplots depict the mean-centred efficiency scores in the relevant condition (given in z-scores) plotted as a function of the normalised functional connectivity (i.e. scaled to the global mean) for left ATL (red) and right functionally-defined ATL (blue) to the cluster depicted in the figure.

### 3.6. Summary of results

We found that higher connectivity between left and right ATL is associated with better categorisation of landmarks, especially for the visual modality. There was a main effect reflecting an association between stronger connectivity and better performance on the picture-based landmark condition, a significant effect of modality for landmarks and an interaction between category and modality that all reflected the same pattern. We summarise the spatial distribution of these effects and their overlap in Figure 7. Distinct from these effects, greater right than left hemisphere connectivity to occipital pole was associated with better performance on landmarks, regardless of modality.

**Figure 7.**
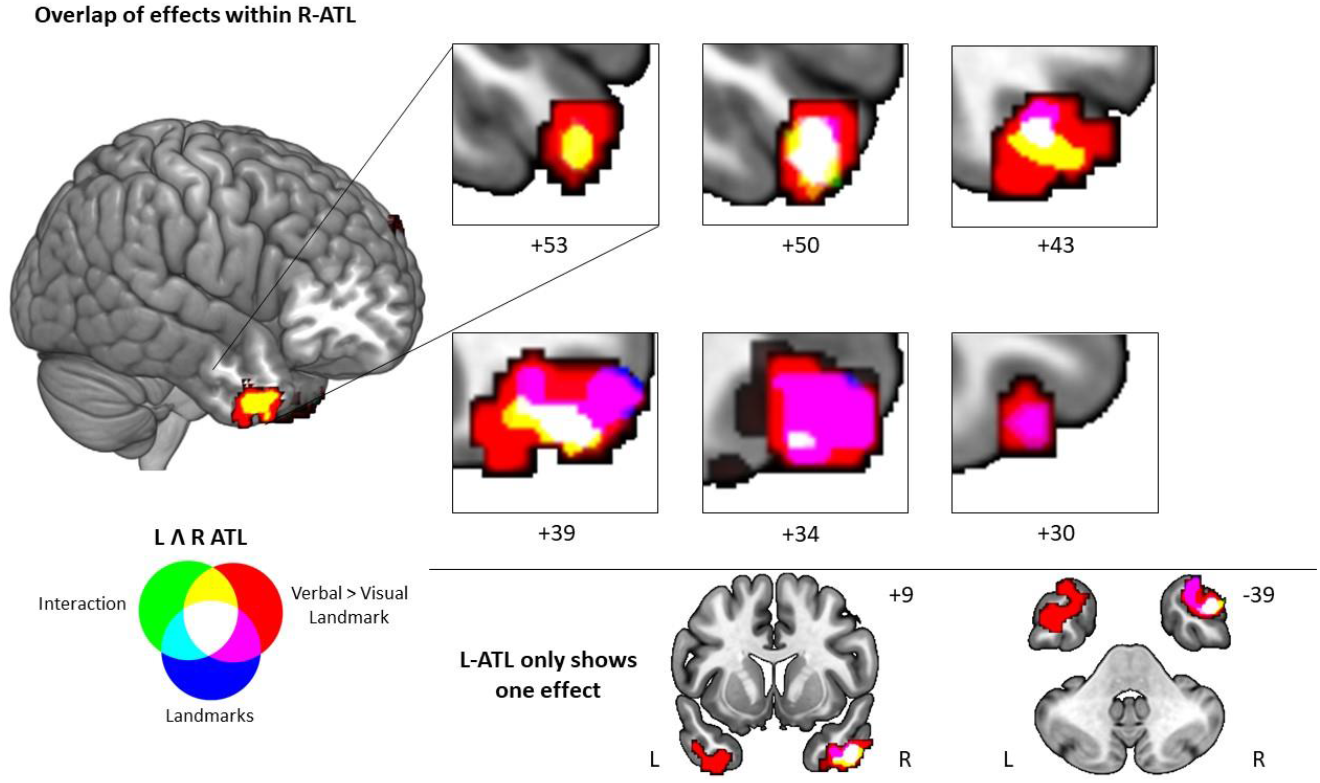
Results of our analysis that fell within left and right ATL (centred on the temporal pole); all of these effects relate to better processing of landmarks, especially in the visual modality. Top panel: Landmarks results that fell in the right temporal lobe and sagittal slices that highlight their topography. Bottom panel: Selected coronal and axial slices that allow comparison between the only effect observed in left ATL with the ones observed in right ATL.

## 4. Discussion

This study examined the relationship between individual differences in the intrinsic connectivity of left and right ATL and performance on semantic tasks that involved (i) different modalities of presentation (pictures vs. words) and (ii) knowledge of specific people vs. landmarks. Previous work has suggested that while the functions of left and right ATL are more similar than they are different, there is some subtle hemispheric specialisation – right ATL might be more important for non-verbal tasks and/or for social concepts, whereas left ATL might play a greater role in written words/naming (Rice et al., 2015b). We used two ATL seeds derived from functionally-defined peaks for left and right ATL, and identified two key behavioural associations with intrinsic connectivity. (i) We found that stronger connectivity from right ATL relative to left to a region in medial / polar occipital cortex was associated with more efficient retrieval of semantic information about famous landmarks (such as determining whether the Eiffel Tower is European), regardless of the modality of presentation. (ii) We also found that when left and right ATL showed strong connectivity to each other (i.e., the system was strongly bilateral), participants were more efficient at retrieving information about landmarks when these items were presented as pictures instead of words.

The first of these key findings -- that efficient categorisation of landmarks is linked to stronger intrinsic connectivity between right ATL and medial regions near the occipital pole -- is consistent with a key role for vision in supporting this category. Landmarks activate two distinct visual processing streams: (i) a dorsal stream that extends from early visual regions into parietal areas and retrosplenial cortex/precuneus, and (ii) a ventral stream originating in the same visual regions, expanding into ventral temporal regions associated with object processing and landmark recognition, and reaching hippocampus, parahippocampal gyrus, fusiform gyrus and ventral anterior temporal cortex (Aguirre and D’Esposito, 1999; Spiers and Maguire, 2007, 2004; Ungerleider, 1982; Yoder et al., 2011). Previous research motivated by the hub-and-spokes account of conceptual knowledge (Lambon Ralph et al., 2017; Patterson et al., 2007) has shown that the dynamic interaction of ventral ATL with medial occipital cortex is essential to the representation and retrieval of visual concepts, even when these are accessed via words (Chiou et al., 2018; Chiou and Lambon Ralph, 2019). In human navigation, landmarks are predominantly visual (although some landmarks, such as a waterfall, may be supplemented by sound). The finding that stronger intrinsic connectivity between right ATL and medial visual cortex is associated with more efficient semantic retrieval for landmarks, regardless of modality, adds to this body of evidence. Right ATL has stronger intrinsic connectivity with medial visual cortex than left ATL (Gonzalez Alam et al., 2019), and participants who show this pattern more strongly show better semantic retrieval for landmarks, perhaps because this category requires strong visuo-spatial imagery.

The second of our findings suggests that there are some functional benefits resulting from strong intrinsic connectivity between the two ATLs. This pattern might be expected for a bilateral semantic representation system: patients with bilateral ATL atrophy who have semantic dementia show more substantial semantic deficits than patients with unilateral ATL lesions following resection for temporal lobe epilepsy (Lambon Ralph et al., 2012; Rice et al., 2018a), perhaps because the two ATLs show a high degree of connectivity (Gonzalez Alam et al., 2019) and consequently the semantic store is only partially divided between left and right hemispheres. This neuropsychological data is accommodated by a model of ATL with strong bilateral connections, as well as somewhat distinct connections from left and right ATL to other brain regions (Schapiro et al., 2013). However, this benefit of bilateral connections between the two ATLs was shown in the current study to be unequal across tasks. Strong cross-hemispheric connectivity particularly benefits tasks which probe knowledge of places and that also use pictorial inputs – perhaps because, in these circumstances, right-lateralised visual-spatial representations (Liu et al., 2009; Stevens et al., 2012) need to be integrated with a left-lateralised semantic retrieval network (including IFG and posterior temporal regions), shown to activate more strongly to semantic decisions about landmarks than people in an on-line fMRI study employing the same tasks (Rice et al., 2018b). Figure 8 shows the neural network engaged by the contrast of landmarks vs. people in the Rice et al. (2018b) study, and its similarity with regions showing stronger intrinsic connectivity to left than right ATL (Gonzalez Alam et al., 2019 and present study). The contrast of Landmarks > People in Rice’s study shows a positive correlation with the left > right ATL connectivity map (r = .21) and a negative correlation with the right > left connectivity map (r = −.23). These correlations are significantly different (Fisher’s r-to-z; z = 1.78, p < .001), suggesting that landmarks may be more reliant on regions more strongly connected to left ATL. This network includes regions of bilateral fusiform and parahippocampal cortex thought to be critical for scene processing (Hodgetts et al., 2016). Participants with more efficient retrieval of semantic information for pictures of landmarks may integrate this network, biased towards left ATL, with right ATL. According to the graded hub account, each ATL receives its strongest inputs from proximal regions within the same hemisphere, and consequently visual-spatial and stimulus-driven attention networks which are right-lateralised (Liu et al., 2009) in posterior temporal and parietal cortex might have privileged access to the right ATL (Gonzalez Alam et al., 2019). One yet untested hypothesis emerging from this analysis is the possibility that patients with semantic dementia might have more severe difficulties in retrieving conceptual information from visual landmark pictures (compared with the categorisation of famous faces and names, and the names of landmarks), reflecting their highly bilateral atrophy.

**Figure 8.**
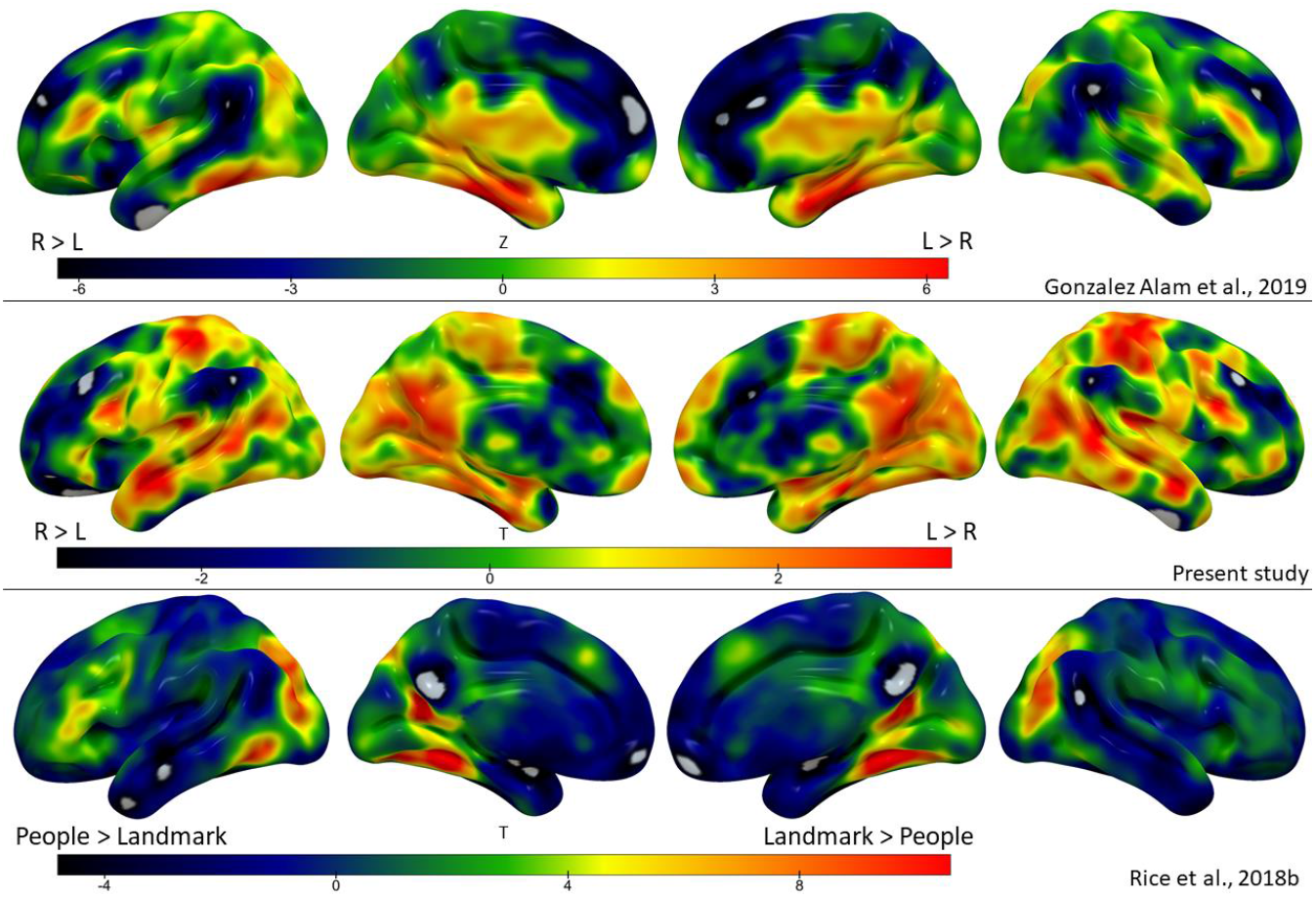
Relationship between Rice et al. (2018b) task-based fMRI results with differential intrinsic connectivity across left and right anterior temporal lobes in Gonzalez Alam et al., 2019 and the present study. The top and middle panels depict the left > right ATL connectivity contrast in Gonzalez Alam et al., 2019 and present study respectively, while the bottom panel shows the Landmark > People contrast in Rice et al., 2018b. The correlations were performed using ‘fslcc’, part of the FSL suite (Jenkinson et al., 2012; Smith et al., 2004). All maps are unthresholded.

In conclusion, this study shows that individual differences in intrinsic connectivity of left and right ATL are associated with effects of category and modality on semantic efficiency in the processing of landmarks. These effects can be interpreted in terms of graded differences in the strengths of inputs from ‘spoke’ regions, such as regions of visual cortex, to a bilateral yet partially segregated semantic ‘hub’, encompassing left and right ATL.

## Acknowledgements

We would like to thank the students that helped with data collection: Melinda Bjolseth, Sophie Colgan, Lucy Milne, Clara Scatola, Nikita Schaap, George Shone, Molly Sibson-Flood, Dragos Teodorov and Molly Wilson.

## Supplementary Materials

### 1. Single seed results

We carried out behavioural regressions on the intrinsic connectivity of left ATL and right ATL separately. The results of these single seed analyses fall within the effects described in the main text for the combined (left plus right ATL) and differential (left versus right ATL) analyses. Left ATL showed two significant results, presented in Figure S1: participants with stronger connectivity from this left ATL seed to a right ATL cluster showed a better performance for visual relative to verbal landmarks; we also found a category by modality interaction in the same region.

**Figure S1.**
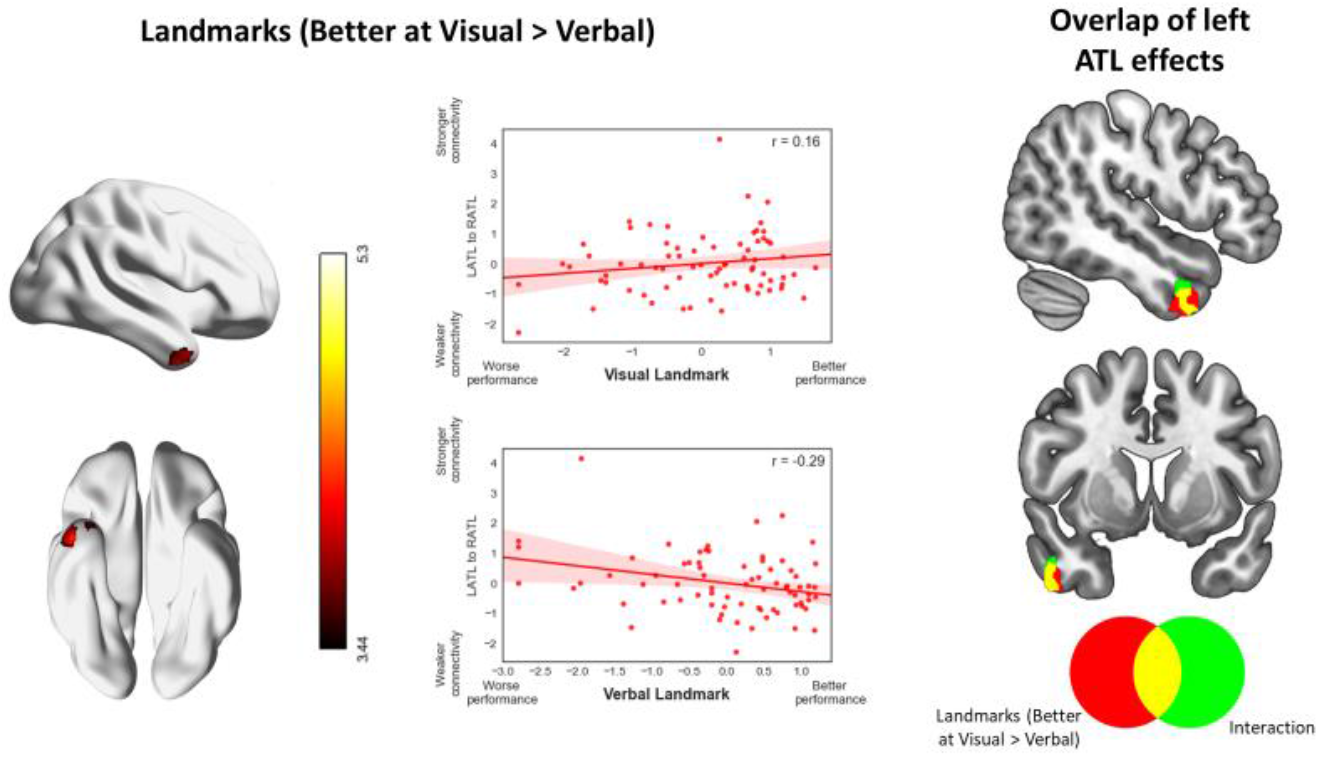
The connectivity between a right temporal pole cluster and the left ATL seed was associated with better performance at visual relative to verbal landmark judgements (Z=3.1, p=.05, Bonferroni-corrected for 4 models). The scatterplots depict the mean-centred efficiency scores (given in z-scores), plotted as a function of the normalised functional connectivity (i.e. scaled to the global mean) between the right temporal pole cluster depicted in the figure and the left ATL seed. This cluster overlapped with an interaction effect, as depicted in the right column of the figure.

Right ATL also showed two significant results: participants with stronger connectivity from the right ATL seed to a more anterior right temporal pole cluster were better at visual relative to verbal landmark judgements, as well as generally better at visual landmarks. These results are shown in Figure S2.

**Figure S2.**
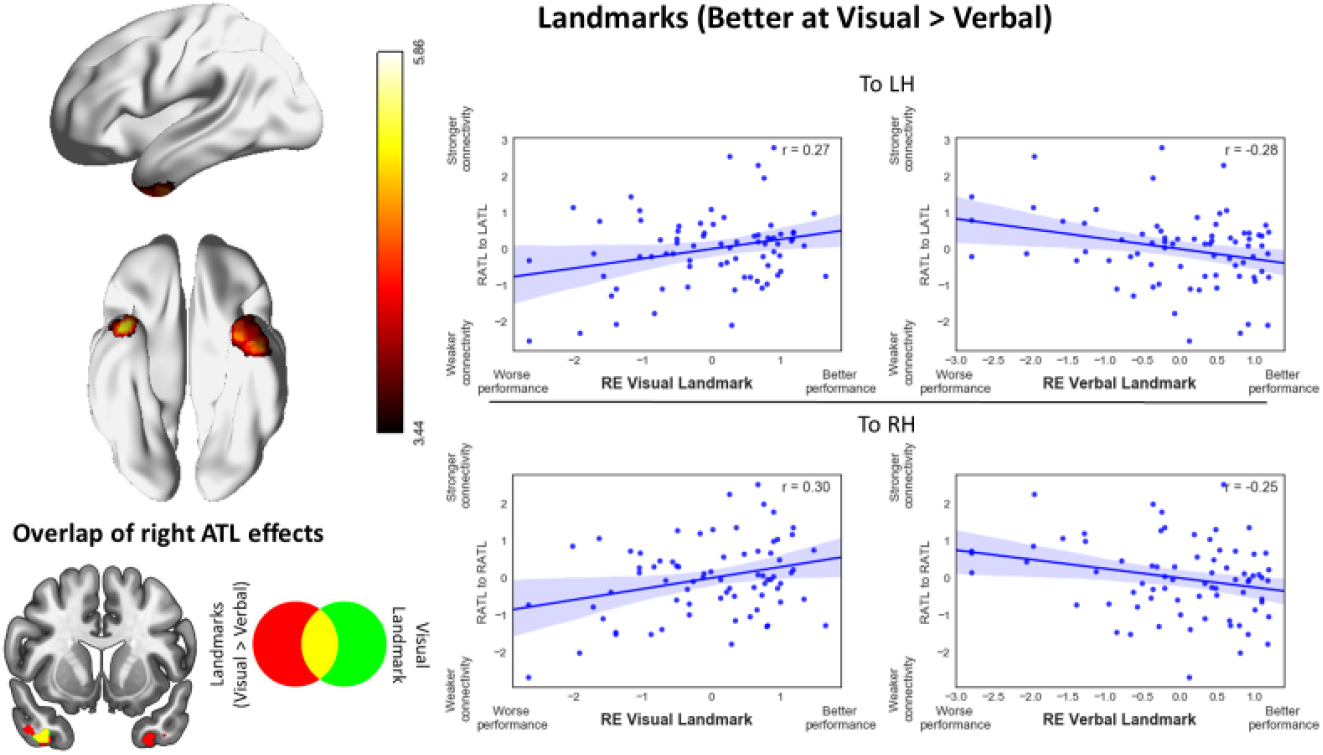
The connectivity between bilateral ATL clusters and the right ATL seed was associated with better performance at visual relative to verbal landmark judgements (Z=3.1, p=.05, Bonferroni-corrected for 4 models). The scatterplots depict the mean-centred efficiency scores (given in z-scores), plotted as a function of the normalised functional connectivity (i.e. scaled to the global mean) between the left and right ATL clusters depicted in the figure and the left and right ATL seeds. The right ATL cluster overlapped with the main effect of being good at visual landmarks, as depicted in the bottom left section of the figure.

### 2. Control analysis with a sign-flipped homotopic seed

#### Rationale

In the main analysis for this study, we used left and right ventral ATL sites identified by Rice et al. (2018) which showed peak activation for semantic > non-semantic contrasts across eight studies using distortion-corrected fMRI (MNI coordinates −41, −15, −31). As these sites were in similar but not identical locations in the two hemispheres, we performed a supplementary control analysis, which compared the left hemisphere functional peak from Rice et al. with a right-hemisphere homotopic seed, generated by flipping the sign of the x coordinate in MNI space from negative to positive. This allowed us to compare the results for functionally-defined seeds with homotopic sites with equivalent MNI coordinates across the two hemispheres (see Figure S3 for a comparison of the location of the homotopic and functional right-hemisphere seeds).

As before, seeds were generated by creating binarised spherical masks with a radius of 3mm. We used the analysis pipeline reported in the main methods, with the exception that the Bonferroni correction was set to correct for 7 multiple comparisons. This accounted for three extra supplementary analyses (the new single right hemisphere seed and both its conjunction and difference analysis contrasting it to left ATL).

**Figure S3.**
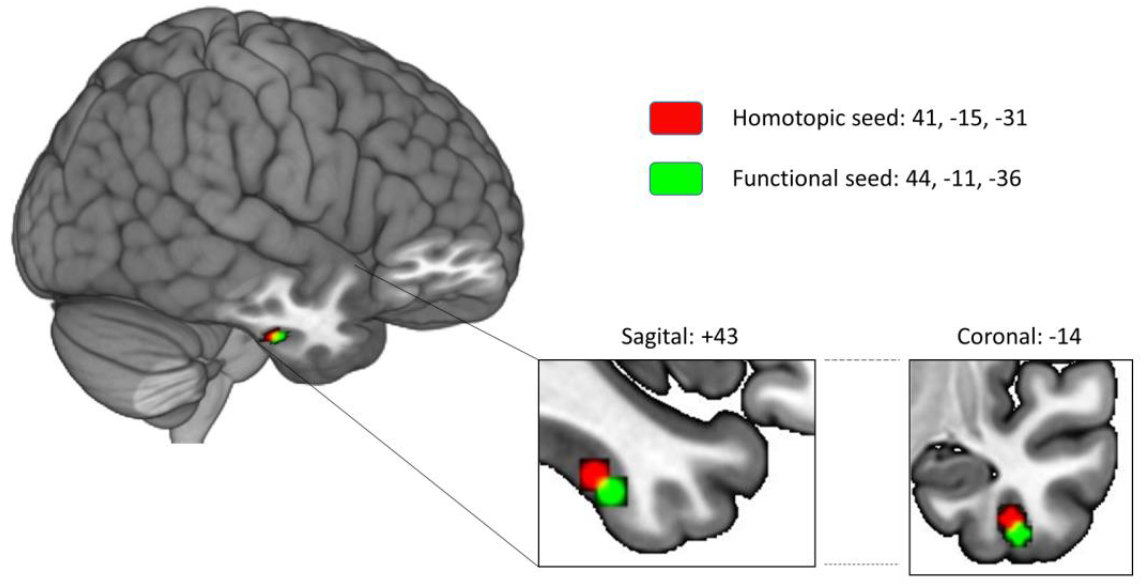
Depiction of the position of the homotopic and functional seeds used for probing right ATL, and their spatial relation to each other.

#### Results

##### a. Resting-state functional connectivity

We examined the mean functional connectivity of left ATL (see main text) and its right homotope using resting-state fMRI. We examined the single seed mean connectivity for each ATL, as well as mean connectivity from both seeds combined, and differential left vs. right connectivity. The results are shown in Figure S4. The RH homotopic seed showed a similar pattern to the LH functional seed: it showed temporal lobe connectivity, albeit less continuous in the LH and more restricted to the ventral temporal lobes, with a separate cluster for angular gyrus that extended more posteriorly than for the left ATL. Unlike left ATL, the RH homotope did not show connectivity to the central sulcus, but there was high intrinsic connectivity with superior frontal gyrus and medial orbitofrontal cortex extending posteriorly to the posterior cingulate cortex; this RH site also showed weak intrinsic connectivity with medial occipital, paracingulate and right insular/orbitofrontal regions (Figure S4, second row). This pattern is similar to that described by Gonzalez Alam et al. (2019). An analysis giving equal weight to both left and right ATL seeds captured this similarity between the maps (Figure S4, third row). Significant differences between the left and right ATL were only observed in the ventral regions centred around our seeds (Figure S4, fourth row). Lastly, a direct comparison between the homotopic and functional right ATL seeds revealed stronger connectivity for the homotopic seed in bilateral LOC, lingual gyrus, posterior cingulate cortex and medial temporal cortex, and stronger connectivity with the functional seed in bilateral vATL, right frontal pole and left temporo-parietal junction (Figure S4, bottom row).

**Figure S4.**
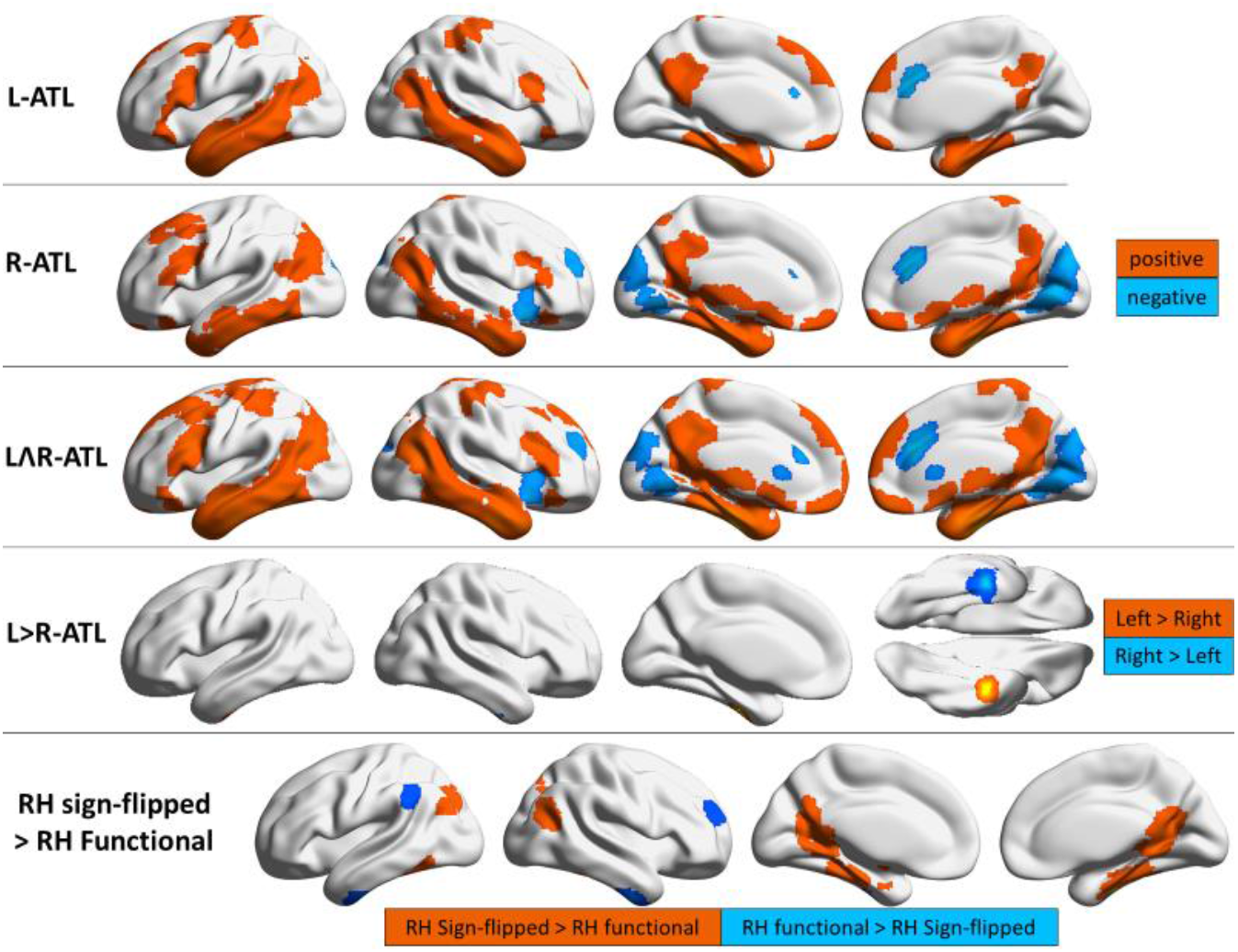
Resting state connectivity for the homotopic peak in right ATL, its common and differential connectivity with functionally-defined left ATL (from Rice et al., 2018), and a comparison of the functionally-defined right ATL with the right hemisphere seed determined through sign-flipping. For the single seeds and mean, the warm and cool colours represent the positive and negative patterns of connectivity respectively, while for the difference analysis, the warm and cool colours represent left and right connectivity respectively. In the last row, stronger connectivity for the homotopic seed is represented by warm colours while stronger connectivity for the functional seed is depicted in cool colours. The group maps are thresholded at z = 3.1, p = 0.05.

##### b. Differential ATL connectivity between hemispheres and associations with behaviour

Our key motivation was to establish if the patterns of connectivity identified through whole-brain analysis of the functional seeds (reported in the main text) could still be observed using a homotopic right ATL seed (employing a regions-of-interest analysis). We also present the results of whole-brain analyses employing the homotopic left and right ATL seeds, for completeness.

To establish whether the results from the functionally-defined right ATL seed could be recovered using the sign-flipped homotope, we performed a series of ROI analyses, using the clusters obtained in our main analysis (Figures 4–6) as a mask. We extracted the average connectivity across the voxels in these masks for the left and right ATL (sign-flipped) seeds, scaled to the global mean for each participant using REX software implemented in CONN. A simple linear regression was calculated for each effect, predicting behavioural performance from the connectivity of left and right (sign-flipped) ATL seeds to each relevant cluster obtained using the functionally-defined right ATL seed (i.e. the analyses in main manuscript). For more complex behavioural effects, repeated-measures ANOVA was used to capture interacting effects of modality and category, with the relevant pattern of connectivity included as a predictor.

The results of these analyses replicated the pattern we obtained using the functionally-defined right ATL seed. The regression for the interaction effect (originally presented in Figure 4) was significant (F(1,72) = 9.189, p = .003). The regression for the visual > verbal landmark effect (Figure 5, top row) was significant for both the left cluster (F(1,72) = 6.190, p = .015), with an R^2^ of .079, and its right counterpart (F(1,72) = 6.394, p = .013), with an R^2^ of .083. The regression for the main effect of visual landmarks (Figure 5, bottom row) was also significant (F(1,72) = 6.394, p = .013), with an R^2^ of .083. Finally, the regression for the left > right ATL main effect of landmarks result was significant (F(1,72) = 4.659, p = .034), with an R^2^ of .061. This pattern of results strengthens our confidence that the results reported in the main text do not solely reflect anatomical differences in seed location across left and right hemisphere.

The whole-brain difference analysis contrasting left ATL with the homotopic seed in right ATL revealed three significant results, all of them located in medial occipital lobe. First, we identified two clusters reflecting an association between connectivity and performance that was common to all tasks (Figure S5). Stronger connectivity from the right ATL homotopic seed was associated with better performance in all tasks, while left ATL connectivity was associated with poorer performance. Next, we observed an interaction effect: this benefit for right-lateralised connectivity was stronger for people’s names than for other conditions (Figure S6). Finally, there was a significant effect of modality for the categorisation of people (Figure S7). Better verbal than visual categorisation of people was associated with differential right versus left ATL connectivity to a relatively large bilateral occipital cluster. The scatterplots show that participants with stronger connectivity to occipital cortex from the right homotopic relative to the left ATL seed had better performance for people’s names.

**Figure S5.**
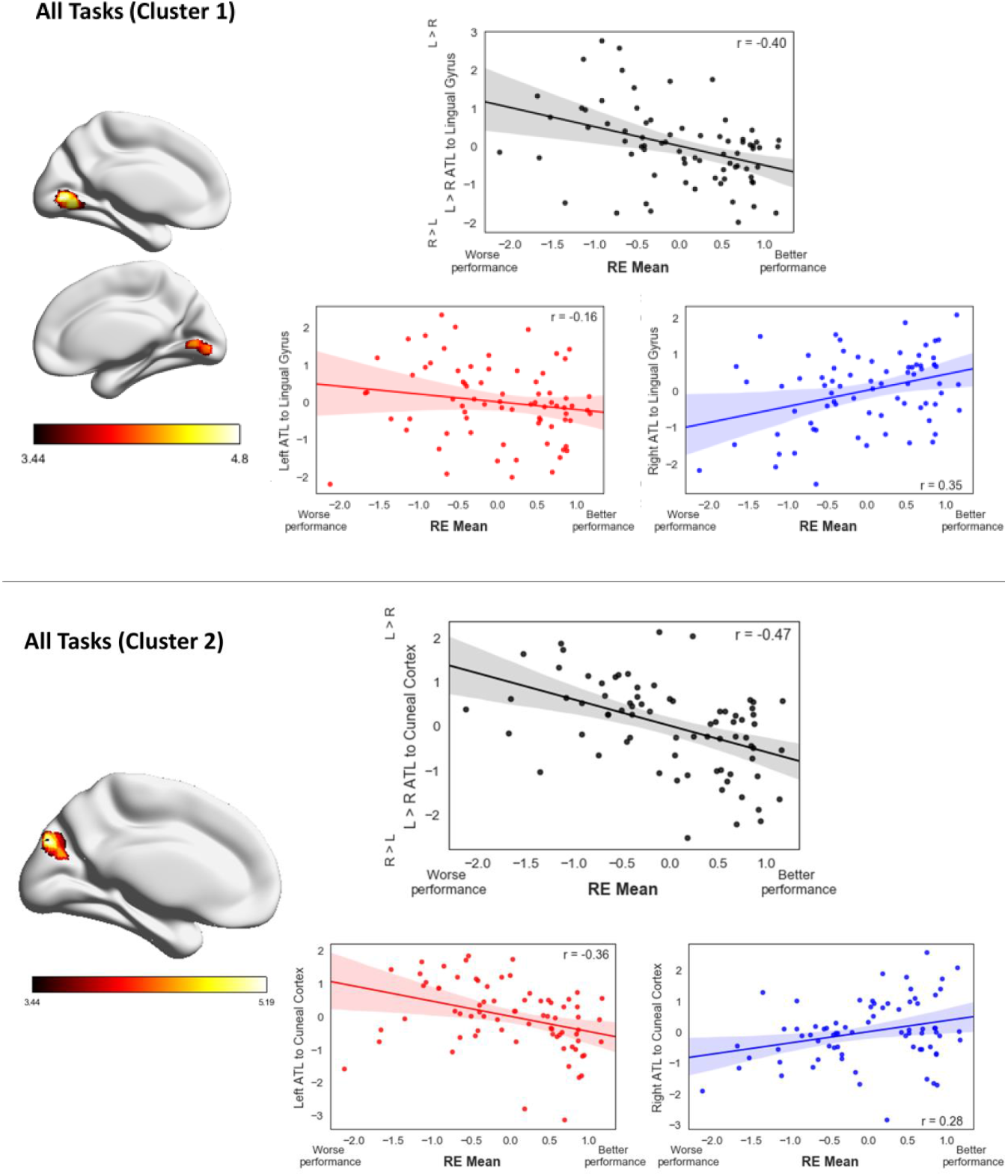
Top panel: A ventral occipital cluster whose differential connectivity to the right homotopic vs. left ATL seed was significantly associated with being good at all tasks. Bottom panel: A dorsal occipital cluster showing the same pattern (results are thresholded at Z=3.1, p=.05, Bonferroni-corrected for 7 models). The scatterplots depict mean-centred efficiency scores averaged across all tasks (given in z-scores), plotted against the normalised functional connectivity (i.e. scaled to the global mean) for differential left versus right ATL (black), left ATL (red) and right homotopic ATL (blue) to the clusters depicted in the figure.

**Figure S6.**
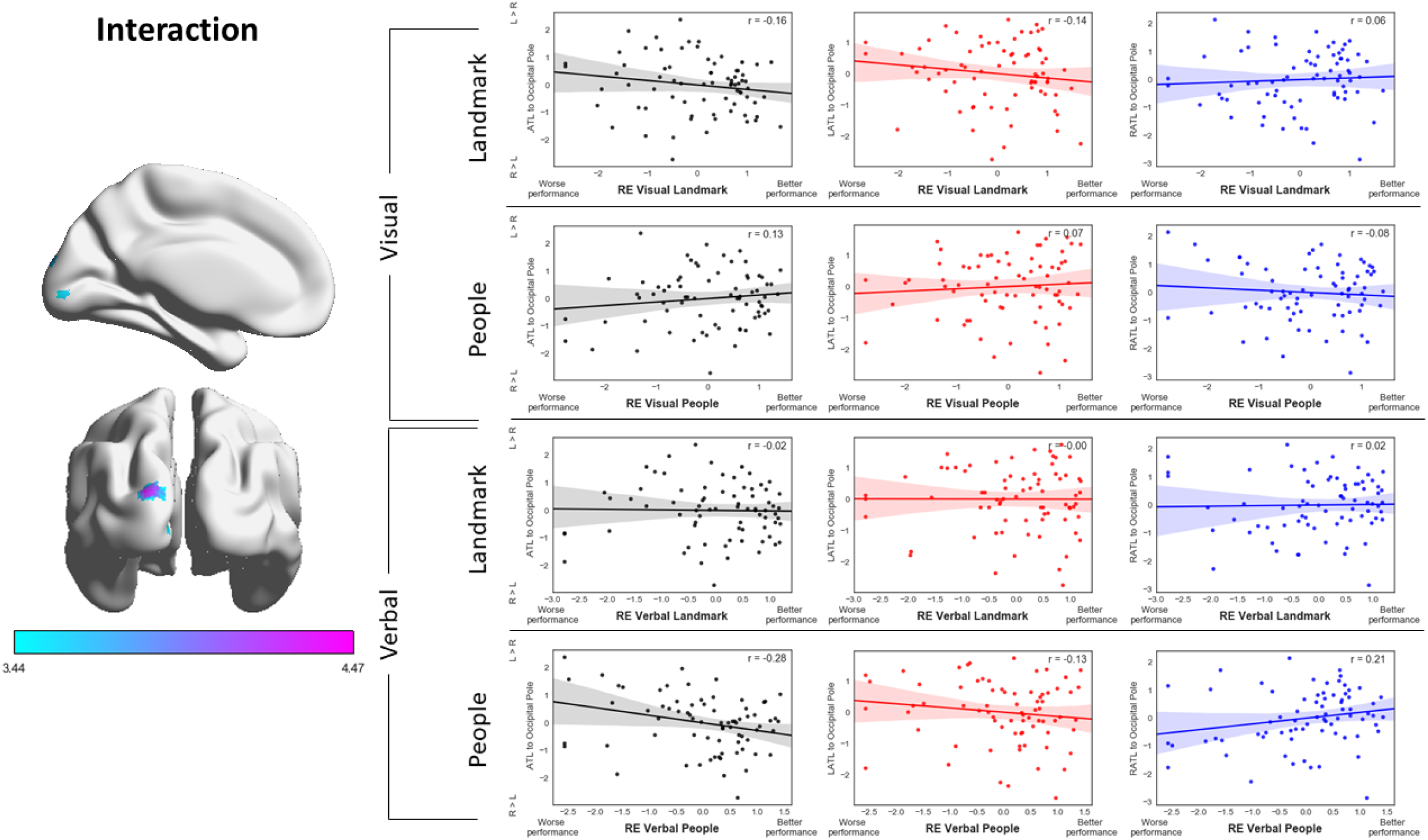
A polar occipital cluster whose differential connectivity to left ATL vs. right homotopic ATL was significantly negatively associated with a modality by category interaction (Z = 3.1, p = .05, Bonferroni-corrected for 7 models). The scatterplots depict the mean-centred efficiency scores in the relevant condition (given in z-scores), plotted as a function of the normalised functional connectivity (i.e. scaled to the global mean) for differential left versus right ATL (black), left ATL (red) and right ATL (blue) to the cluster depicted in the figure.

**Figure S7.**
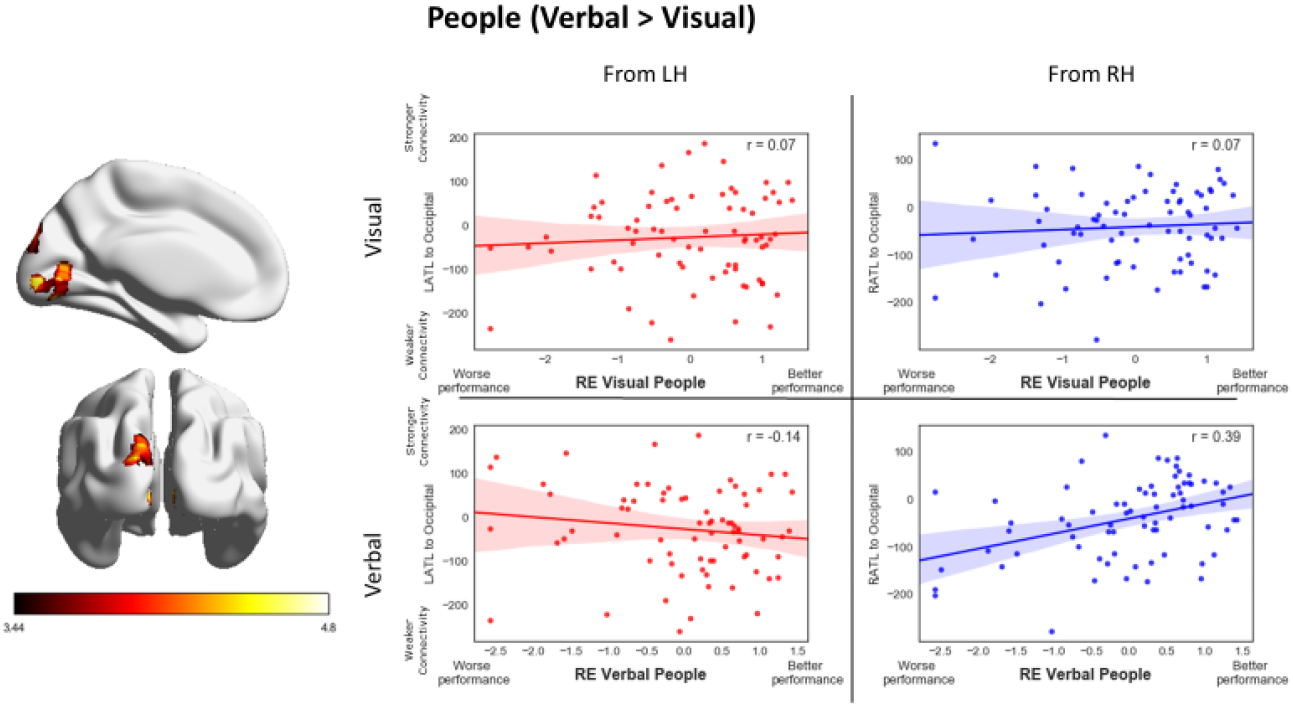
A bilateral occipital cluster whose differential connectivity to left ATL vs. the right homotopic ATL was significantly associated with being better at verbal than visual judgements of people (Z=3.1, p=.05, Bonferroni-corrected for 7 models). The scatterplots depict the mean-centred efficiency scores in the relevant condition (given in z-scores), plotted as a function of the normalised functional connectivity (i.e. scaled to the global mean) for left ATL (red) and right ATL (blue) to the cluster depicted in the figure. All units are given in z-scores.

In sum, using homotopic seeds, we found stronger connectivity between right ATL and ventral visual cortex was associated with better performance across all tasks, although there were some nuances: stronger connectivity between right ATL and a ventral occipital cluster was associated with better semantic performance, while stronger connectivity between left ATL and a more dorsal occipital cluster was associated with poorer performance. There was a significant interaction between category and modality, and a modality effect for the people category, reflecting a bigger benefit of right ATL connectivity for the categorisation of people’s names. Distinct from these effects, greater right than left hemisphere connectivity to occipital pole from functionally-defined right ATL (as reported in the main results section) was associated with better performance on landmarks, regardless of modality. We present a summary of the spatial distribution of the occipital results and their overlap in Figure S8.

**Figure S8.**
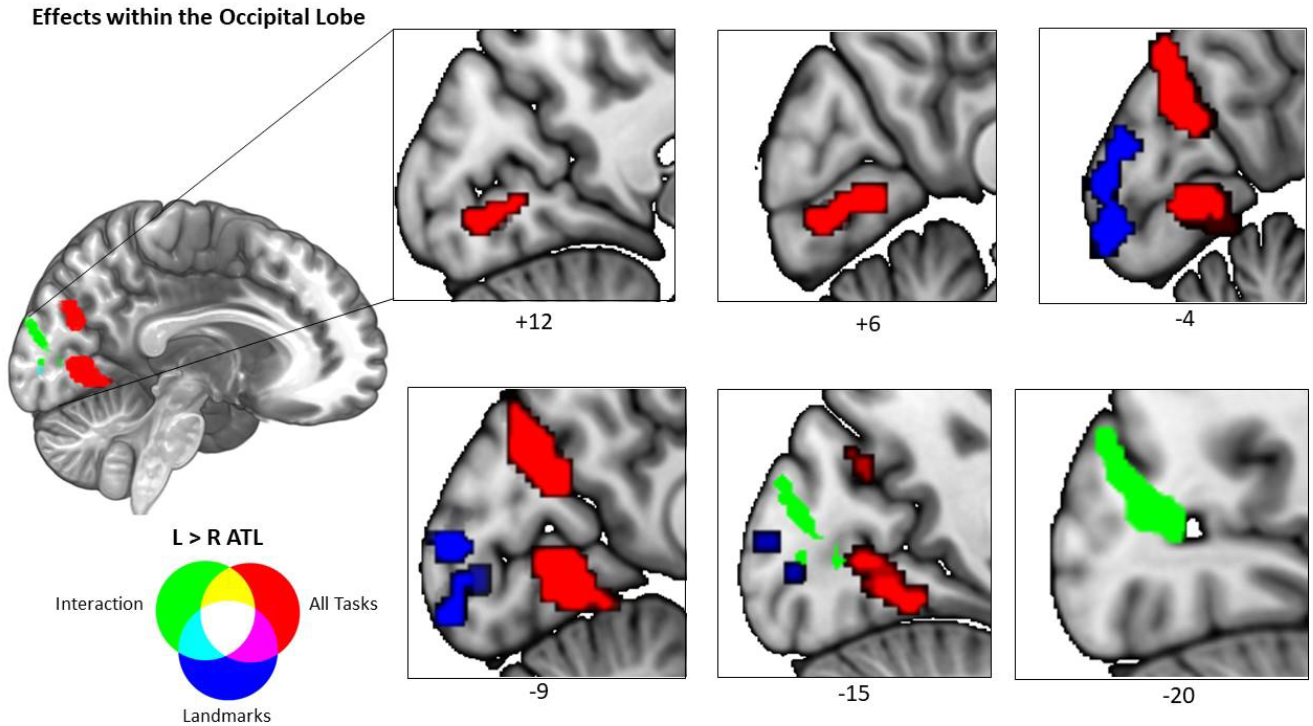
Results of differential analyses that fell in the occipital lobe, depicted in selected slices to highlight their relative locations. These results capture two general effects: being differentially good or bad at tasks as a function of right or left ATL connectivity to these clusters, and an effect of being good at landmark judgements regardless of modality. Taken together, these clusters span a large part of the medial occipital lobe and show minimal overlap. The landmarks result stems from the left > right functional seed analysis, whilst the interaction and all tasks effects come from the left > right homotopic seed analysis.

#### Discussion

The key finding of this supplementary analysis was that stronger intrinsic connectivity from homotopic right ATL to ventromedial visual cortex was associated with more efficient categorisation, especially better retrieval of conceptual information about people from their names. Since connectivity to visual cortex is stronger for right than left ATL (Gonzalez Alam et al., 2019), an exaggeration of the typical pattern of asymmetric connectivity is beneficial to performance. Yet the association between right ATL-to-visual connectivity and an advantage for people’s names is counterintuitive, given written words are more likely to activate left than right ATL (Rice et al., 2015b). A role of the right ATL in knowledge about people, however, is anticipated by previous studies (Gainotti, 2007a, 2007b; Olson et al., 2013; Rice et al., 2015b; Snowden et al., 2004). Using the same paradigm as in the current study, Rice et al. (2018) found a cluster in right vATL for people versus landmark categorisation (see Figure 2, top row in their paper). In addition, the wider network of regions activated by this contrast overlaps with brain regions that show stronger connectivity to right than left ATL (see Figure 8; main manuscript). Rice et al. (2018) also reported a ventromedial visual cluster for the contrast of social words vs. non-social abstract terms. These findings taken together are consistent with the view that right ATL does not specifically support visual semantic cognition; instead connectivity from right ATL to ventromedial visual cortex may play an important role in social cognition, as opposed to face recognition per se. Accessing conceptual information about people from their names as opposed to their faces might entail more visual imagery, thought to be supported by medial aspects of visual cortex (Kosslyn et al., 1999, 1995). Overall, our findings do not fit well with the hypothesis that left ATL differentially supports verbal tasks, while right ATL supports picture-based tasks. Our results are more compatible with the view that right ATL plays a central role in knowledge of people, and that integration of conceptual processes with visual cortex particularly supports the ability to recall someone from their name.

This supplementary analysis focussed on comparing two right ATL seeds: (i) a functional peak derived from a meta-analysis of semantic task data, and (ii) a homotopic site in the same location as the left ATL peak but sign-flipped to the right hemisphere. These two ATL seeds yielded different behavioural associations (discussed in the main text and this supplementary analysis) – strong intrinsic connectivity from the right ATL functional seed to left ATL was associated with more efficient conceptual retrieval about landmarks, while strong intrinsic connectivity from the right ATL homotopic seed to ventromedial visual cortex was associated with more efficient conceptual retrieval from people’s names. This difference between seeds requires further exploration but might reflect the complex functional organisation of ATL (Jackson et al., 2017, 2016; Rice et al., 2015a). This complex organisation can be seen when the intrinsic connectivity of the right ATL homotopic seed is compared with the functional seed (Figure S4, bottom row); even the minimal distance between these seeds (see Supplementary Figure S3) was enough to engage slightly different networks, with the homotopic (more posterior) seed showing greater connectivity with parts of the ventral visual stream and occipital cortex, compatible with the idea of functional change across a graded hub (Bajada et al., 2019; Binney et al., 2012; Rice et al., 2015a).

